# LncRNA *Spehd* regulates hematopoietic stem cells and progenitors and is required for multilineage differentiation

**DOI:** 10.1101/340034

**Authors:** M Joaquina Delás, Benjamin T Jackson, Tatjana Kovacevic, Silvia Vangelisti, Ester Munera Maravilla, Sophia A Wild, Eva Maria Stork, Nicolas Erard, Simon RV Knott, Gregory J Hannon

## Abstract

Long non-coding RNAs (lncRNAs) show patterns of tissue- and cell-type-specific expression that are very similar to those of protein coding genes and consequently have the potential to control stem and progenitor cell fate decisions along a differentiation trajectory. To understand the roles that lncRNAs might play in hematopoiesis, we selected a subset of mouse lncRNAs with potentially relevant expression patterns and refined our candidate list using evidence of conserved expression in human blood lineages. For each candidate, we assessed its possible role in hematopoietic differentiation in vivo using competitive transplantation. Our studies identified two lncRNAs that were required for hematopoiesis. One of these, Spehd, showed defective multi-lineage differentiation, and its silencing yielded common myeloid progenitors deficient in their oxidative phosphorylation pathway. This effort not only suggests that lncRNAs can contribute to differentiation decisions during hematopoiesis but also provides a path toward the identification of functional lncRNAs in other differentiation hierarchies.

## Introduction

Long noncoding RNAs (lncRNAs) can function as regulators of cell fate via a number of different mechanisms, ranging from regulation of gene expression to effects on mRNA and protein stability (Delás and Hannon, 2017). While there are tens to hundreds of thousands of lncRNAs annotated in different genomes (Melé and Rinn, 2016), the challenge of identifying those lncRNAs that regulate a particular process is still significant. With the breadth of information the field has produced on hematopoietic stem cell (HSC) differentiation, the hematopoietic system is the perfect model to investigate how lncRNAs regulate differentiation in general and how their roles fit within existing paradigms (Orkin and Zon, 2008). The power of this system specifically hinges on the ability to perform *in vivo* bone marrow reconstitutions, which provide the ultimate proof of biological relevance.

Regulation of cell fate transitions during hematopoiesis has been studied at many different levels. A number of transcription factors have been shown to be essential for the various steps of hematopoietic differentiation (Orkin and Zon, 2008), and they are also often positive regulators of their own transcription, forming a highly dynamic transcription factor network (Schütte et al., 2016). This tight regulation of gene expression is also highly dependent on additional transcriptional control mechanisms. DNA methylation changes are associated with regulatory areas where lineage-specific transcription factors bind (Bock et al., 2012; Hodges et al., 2011), and, consequently, the different DNA methyl transferases have been shown to play essential roles in hematopoietic differentiation (Challen et al., 2011; 2014; Trowbridge et al., 2009). Hematopoiesis is also highly sensitive to alterations to chromatin modifiers, both during normal differentiation (Kerenyi et al., 2013) and during malignant transformation (reviewed in (D. Hu and Shilatifard, 2016)).

Post-transcriptional regulation through microRNAs has also been described to contribute to decisions during hematopoiesis. *MicroRNA-126* promotes HSC quiescence, and its knockdown leads to HSC expansion (Lechman et al., 2012) and *in vivo* to hematological malignancies (Lechman et al., 2016; Nucera et al., 2016). This microRNA and its functions are conserved in both mouse and human, as is *miR-223* (Fukao et al., 2007), required for granulocyte differentiation (Johnnidis et al., 2008).

The function of several lncRNAs has been addressed in *in vitro* models of hematopoietic differentiation, such as granulocyte differentiation (Zhang et al., 2009), eosinophil differentiation (Wagner et al., 2007), and erythropoiesis (W. Hu et al., 2011). Global analysis of annotated lncRNAs has also revealed that their expression is regulated in early stem cell populations (Cabezas-Wallscheid et al., 2014). Since the current GENCODE annotation for lncRNAs is mostly based on easy-to-culture cell lines or whole organisms, it lacks many of the cell-type-specific hematopoietic transcripts. To circumvent this, some groups have assembled annotations for subsets of the hematopoietic lineage or for some of the differentiation models mentioned above (Alvarez-Dominguez et al., 2014; Luo et al., 2015; Paralkar et al., 2014). We recently sought to produce a robust annotation that encompassed cell types from HSCs to differentiated cells, both myeloid and lymphoid lineages, as well as blood cancers (Delás et al., 2017). In our first proof-of-concept study, we used this resource to characterize lncRNAs required for acute myeloid leukemia.

Here we focused on characterizing lncRNAs involved in the earliest choices the HSC has to make: selfrenewal or commitment to a lineage. In order to address this question, we devised an experimental strategy whereby long-term reconstituting stem cells could be transduced *in vitro* with shRNAs targeting lncRNAs and then transplanted to uncover lncRNA dependencies *in vivo*. Moreover, since many aspects of hematopoiesis are conserved between mouse and human (Shay et al., 2013), we reasoned that if we could identify lncRNAs with syntenic conservation and conserved expression in human, we could enrich for functional potential.

## Results

### Combination of differential expression with syntenic conservation and conserved expression to enrich functional lncRNAs

To maximize the likelihood of identifying lncRNAs functionally required for hematopoietic stem cell differentiation and/or stem cell self-renewal, we designed a selection strategy to prioritize lncRNAs that show evidence of conserved differential expression in key hematopoietic progenitors. LncRNAs are not often conserved in sequence, but one can often find a lncRNA in a syntenic position in different genomes (Hezroni et al., 2017).

Our first criterion applied in assembling a candidate list was that a lncRNA would have an expression pattern consistent with its being a regulator of hematopoiesis. We hypothesized that if the expression of a candidate lncRNA was tightly regulated during hematopoietic differentiation, it would be more likely to be involved in regulating cell fate transitions during that differentiation process than would a ubiquitously expressed transcript. We combined lncRNAs with (1) enriched expression in hematopoietic stem cells (HSC), committed myeloid or granulocyte/monocyte (CMP/GMP), or lymphoid progenitors (CLP) when we analyzed differential expression between those cell types (progenitor cell-type enriched), (2) differential expression between myeloid and lymphoid lineages that already displayed primed differential expression at the progenitor level (lineage enriched) and (3) differential down-regulation transcripts during differentiation while already differentially expressed between any of the progenitor/stem populations (down-regulated in differentiation) (Fig. 1A). This generated a list of 295 mouse lncRNAs with suggestive expression patterns (Fig. S1A). Since our functional assay relied on gene silencing in transplanted, modified HSCs, we decided not to include lncRNAs that were downregulated between progenitors and differentiated cell types, as we reasoned that these would require an enforced expression assay to reveal impacts during differentiation.

**Figure 1.**
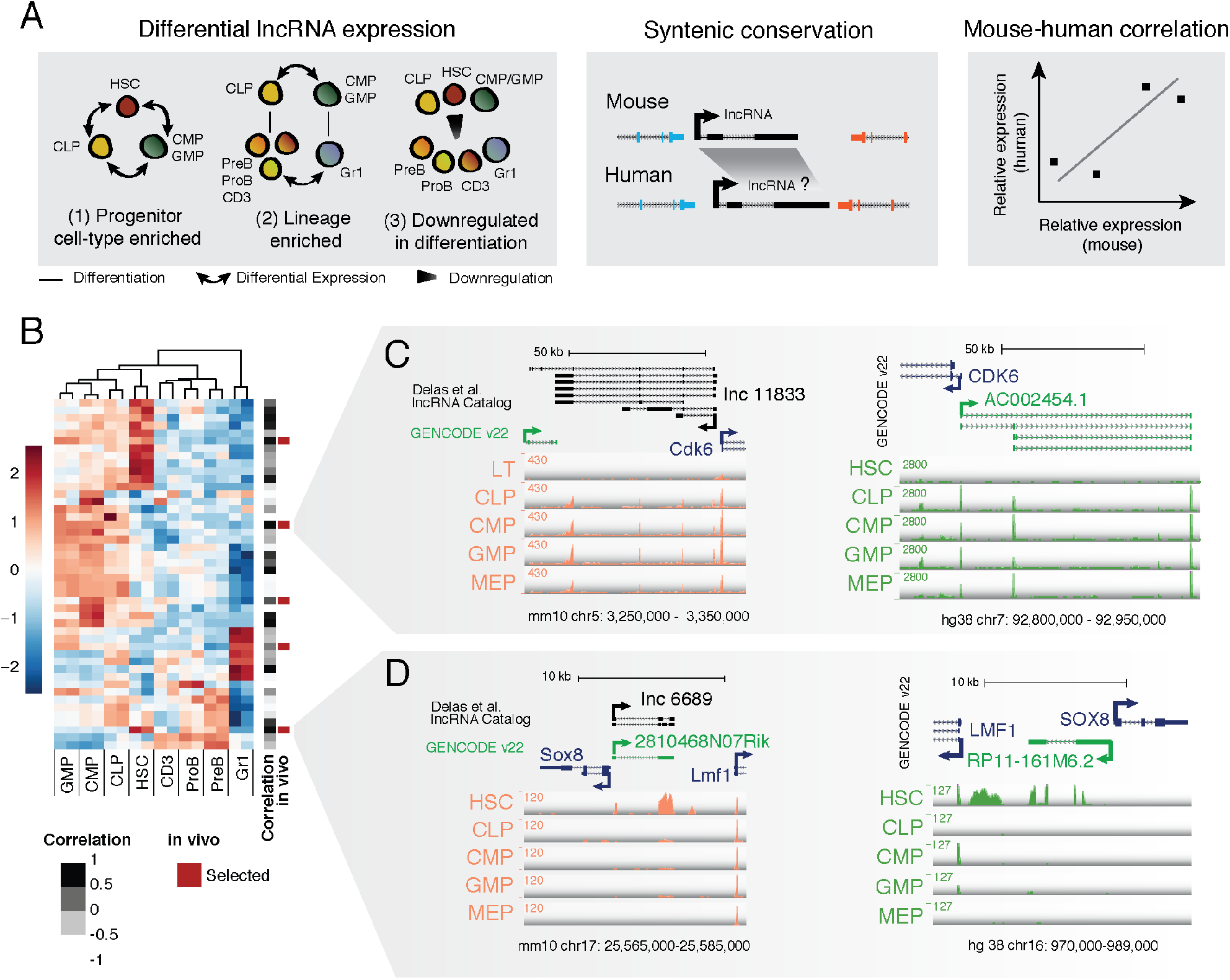
An approach to enrich potentially functional lncRNAs. (A) Pipeline overview from the expression patterns selected to mouse-human expression correlation (B) Expression heat map of the 46 mouse lncRNAs that have expressed syntelogs in human, indicating their level of expression correlation and the lncRNAs selected for *in vivo* studies. (C-D) Genome browser plots for the mouse (left) and human (right) loci for lnc11833-AC002454.1 (C) and lnc6689-RP11-161M6.2 (D) with expression for a representative replicate of each of the progenitor cell types.

Using the UCSC liftOver utility we found that, in 97 cases, we could identify an annotated lncRNA in a syntenic position in the human genome. We then analyzed the expression of those lncRNAs in human cord blood HSC, CMP, GMP and CLP (Chen et al., 2014), yielding a list of 45 lncRNAs that were expressed in these cell types.

We analyzed the expression correlation for each of these lncRNAs in human and mouse for the aforementioned cell types (Fig. 1B) to prioritize lncRNAs for *in vivo* studies. Due to limitations in lncRNA assemblies and the complex genomic organization of some loci, we assembled our final list of 5 lncRNAs after manual inspection of each genomic locus to ensure that the human expression data was reflective of the lncRNA levels (and not an overlapping gene) and that the lncRNAs selected showed a consistent exon structure across replicates. Additionally, we tried to maintain a representation of the different expression patterns initially selected. *Lncll833*, for example, is divergently transcribed with *Cdk6* and dramatically upregulated in all committed progenitors as compared to stem cells in both mouse and human (Fig. 1C). Given the reported role of Cdk6 in regulating human hematopoietic stem cell quiescence (Laurenti et al., 2015), the location and expression of this lncRNA was appealing and potentially suggestive of an effect in *cis*. Another lncRNA, *lnc6689*, is specifically expressed in stem cells but not in the progenitor cell types (Fig. 1D). While *lnc6689* (annotated in GENCODE as 2810468N07Rik) is also expressed divergently from a neighboring protein-coding gene, *Sox8*, this gene is not expressed in blood progenitors in either human or mouse and does not have any known role in hematopoiesis.

We also identified an already described lncRNA that overlaps with microRNA-223 (Fig. S1B). This microRNA plays a role in myelopoiesis (Fazi et al., 2007; Fukao et al., 2007) and the lncRNA has recently been implicated in acute myeloid leukemia (Mangiavacchi et al., 2016). A complete list of the lncRNAs selected for *in vivo* studies and their coordinates can be found in Table S1 and the corresponding genome browser visualizations in Fig. S1B-D.

### Bone marrow transplantation identifies lncRNAs required for hematopoiesis *in vivo*

To assess the functional importance of our selected lncRNAs *in vivo*, we performed bone marrow reconstitutions, transplanting HSCs transduced with a short hairpin RNA (shRNA) against the lncRNAs. Each short hairpin against the candidate lncRNAs was cloned into a constitutive lentiviral vector where an SFFV promoter drives expression of a green fluorescent protein (zsGreen), a Neomycin resistance gene, and the short hairpin. CD45.2 E-SLAM HSCs were sorted (Kent et al., 2009) and transduced with high titer lentivirus for ~ 20h before injecting them into irradiated recipient CD45.1 animals. The cells were mixed with competitor CD45.1 whole bone marrow prior to injection. Starting at 4 weeks post bone marrow transplant, peripheral blood from recipient animals was analyzed by flow cytometry (Fig. 2A, Fig. S2A). The donor compartment (CD45.2) corresponds to the injected HSCs whereas the CD45.1 compartment includes the competitor cells and any remaining cells from the recipient mouse. Because only a fraction of HSCs transplanted had been transduced, we monitored the zsGreen percentage within the donor compartment over time as a phenotypic readout. If the shRNA-expressing cells (also expressing zsGreen) are less able to repopulate the hematopoietic compartment as compared to the non-transduced counterparts, we would expect a depletion of zsGreen-positive cells over time.

**Figure 2.**
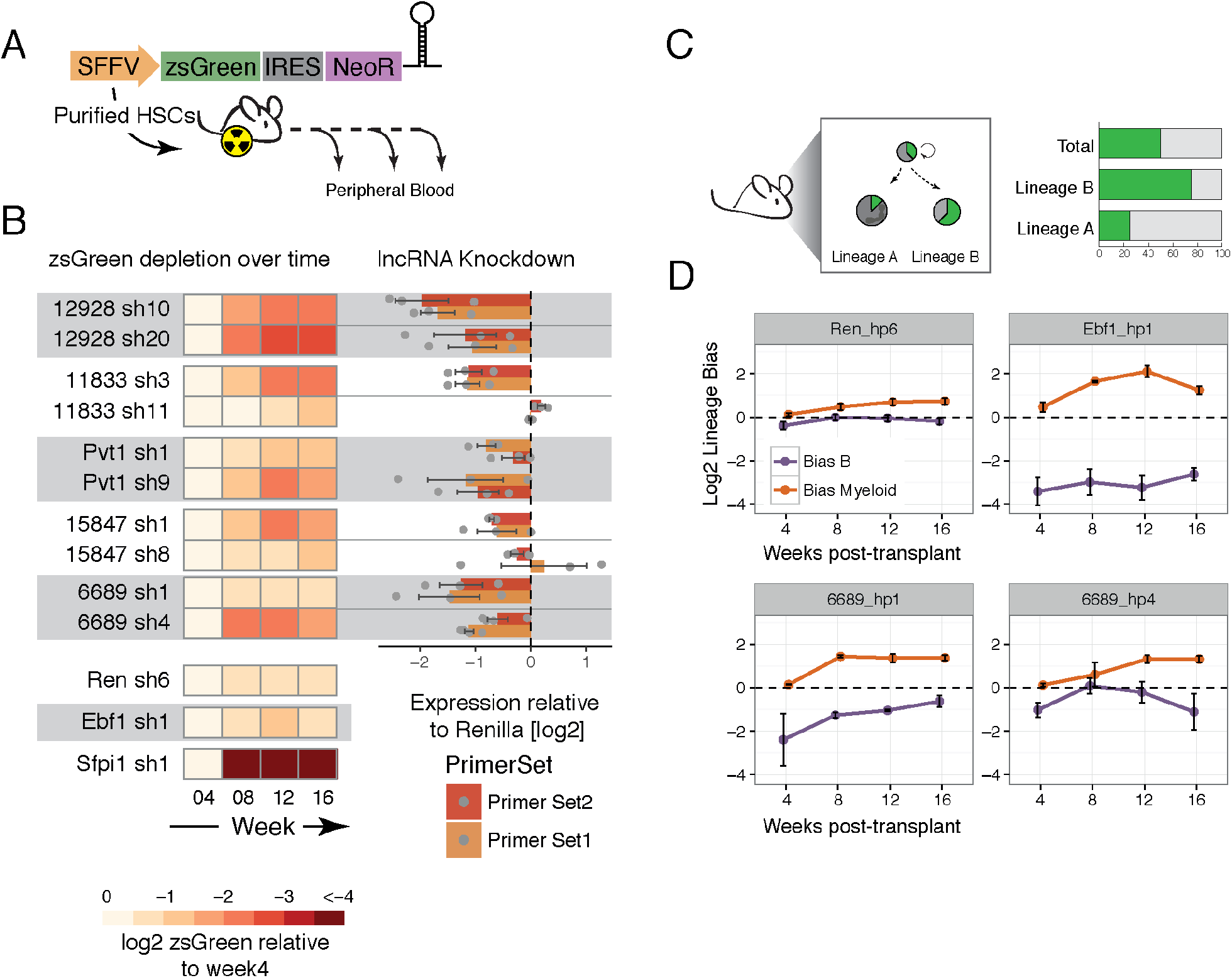
In vivo reconstitution with lncRNA-depleted HSCs identifies lncRNAs involved in overall differentiation or lineage specification. (A) Schematic representation of the vector used and the experimental design. (B) Heat map indicating the average depletion of zsGreen-postiive cells relative to the week 4 measurement (left) and the corresponding level of knock down in a cell line that expresses the corresponding lncRNA (see Experimental Procedures) for each short hairpin assayed in vivo. (C) Schematic representation of the concept of lineage bias analysis. (D) Average lineage bias for the myeloid and B lineages at the different time points in the blood for the indicated knockdowns. Error bars represent s.e.m.

Indeed, we saw a dramatic decrease in the representation of zsGreen-positive cells transduced with an shRNA against positive control *Sfpi1* (also known as *PU1)* as compared to cells expressing a short hairpin against our negative control, *Renilla* luciferase (Fig. 2B). *Sfpi1* is a transcriptional activator that is indispensable for HSC self-renewal as well as commitment to and maturation of myeloid and lymphoid lineages (Iwasaki et al., 2005). We therefore would expect almost no output of differentiated cells from HSCs in which *Sfpi1* was silenced. Of note, because we do not have a robust way of measuring initial HSC infection rates (getting a reliable measurement of the initial HSC infection would require many animals since they are a very rare population), we normalize to the initial blood measurement at four weeks post-transplant. By that point, the percentage of zsGreen-expressing cells could already be greatly reduced if the targeted RNA was required for HSC differentiation or maintenance.

Using this readout, we identified *lnc12928* as our strongest candidate. Using two independent short hairpins, we saw between a 4 and 8-fold average reduction in the relative number of zsGreen-positive cells within the donor compartment during the course of the experiment (16 weeks) (Fig. 2B). To verify that we were indeed affecting the expression of the targeted lncRNAs with our short hairpins, we performed RT-qPCR on cell lines that express each of the targeted lncRNAs (MLL-AF9;NRAS^G12D^ AML (Zuber et al., 2011) for all lncRNAs except for *lnc6689*, for which we use A20 cells). Knockdown measurements were performed after inducing the short hairpin expression for 48h in clonal cell lines selected for uniform zsGreen induction. We performed each experiment with at least 3 independent clonal cell lines for each short hairpin. Using this approach, we confirmed that the levels of *lnc12928* were greatly reduced using both short hairpins and when measured with two independent primer pairs (Fig. 2B).

Interestingly, we noticed that the only short hairpin that successfully silenced *lnc11833* was the only shRNA that gave rise to a depletion of zsGreen-expressing cells in vivo (Fig. 2B). This suggests that *lnc11833* could still be a potentially important lncRNA in hematopoiesis, although additional lncRNA depletion experiments with other tools would be required to validate this observation.

These results show that our HSC transduction followed by transplantation approach is a useful way to identify genes required for hematopoietic reconstitution in general, and is a powerful tool to identify functional long noncoding RNAs.

### HSCs depleted of lncRNA 6689 display a lineage-bias phenotype

lncRNAs could impact hematopoiesis in a variety of ways that would not necessarily lead to changes in the overall representation of zsGreen-positive cells. For example, our lineage control in this assay is Early B cell factor 1 *(Ebf1)*, a gene required for B cell differentiation (Zandi et al., 2008). *Ebf1* knockdown does not result in changes in the relative fraction of zsGreen-positive cells, as compared to week 4 post-transplant (Fig. 2B). However, we observed that within each animal, the percentage of zsGreen-positive cells in the B220-positive compartment was much lower than in the myeloid cell types. To investigate this further, we assessed, within each animal, the percentage of zsGreen-positive cells within the B cell donor compartment relative to the overall zsGreen-positive population derived from the donor (lineage bias), and performed the same analysis for the myeloid lineage (Ly6G+ and/or Cd11b+) (Fig. 2C, Fig. S3B). This showed a clear bias against the B lineage and enrichment of the myeloid compartment for *Ebf1* knockdown (Fig. 2D). *Renilla* shRNAs, in contrast, showed a generally balanced phenotype, where the fraction of zsGreen-positive cells within each lineage mirrored the representation of zsGreen-positive populations in blood overall.

By performing the same lineage-bias analysis, we found that one of our lncRNAs, *lnc6689* had a potential bias against B cells or in favor of the myeloid lineage (Fig. 2D, bottom panels). We see this phenotype with both short hairpins targeting this lncRNA, although the extent of the effect differs. This indicated that each zsGreen-expressing HSC transplanted is producing fewer B220-positive progeny or more myeloid cells. The extent of this phenotype correlated with the degree of knockdown we observed when this lncRNA was targeted in A20 cells (Fig. 2B), suggesting a dose-dependent effect.

*Lnc6689* initially caught our attention because of its striking HSC-specific phenotype (Fig. 1D). However, when we examined the expression of *lnc6689* in more mature cell types, we observed its presence in Pre-and Pro-B cells (Fig. S3A), which means that the lineage-biased phenotype could be mediated either by its functioning in HSCs or via an effect during B cell differentiation.

To establish a strategy where we could address potential cell-type-specific effects, we built an inducible shRNA vector that would allow us to perform bone marrow reconstitutions and only activate short hairpin expression once the hematopoietic system has been repopulated. With this system we could acutely induce a short hairpin and investigate its impact in different cell types or at different time points.

We transduced cells with inducible short hairpins against *lnc6689* or controls, *Renilla* and *Ebf1*, and transplanted them into cKit(w41);CD45.1, a mouse strain that is particularly suitable as a HSC recipient and that only requires sub-lethal irradiation. We allowed the animals to reconstitute and induced the short hairpin by feeding doxycycline-containing food (Fig. 3A). When we looked at the whole bone marrow (after lysing red blood cells) of these animals, we saw a bias against the B220+ compartment for cells in which *Ebf1* was silenced, our lineage control. The extent of the bias increased in magnitude if we administered doxycycline to the animals for 6 days, rather than 2 days (Fig. 3B). Although we also see a bias against the B220+ compartment in the *lnc6689* knockdown, the phenotype is less severe and quite variable, especially with short hairpin 4 (Fig. 3B). This knockdown also showed a milder phenotype in the blood (Fig. 2D) and a lower knockdown efficiency (Fig. 2B). Additional experimental efforts will be required to further validate this phenotype and their role in either stem cells or the B cell compartment directly.

**Figure 3.**
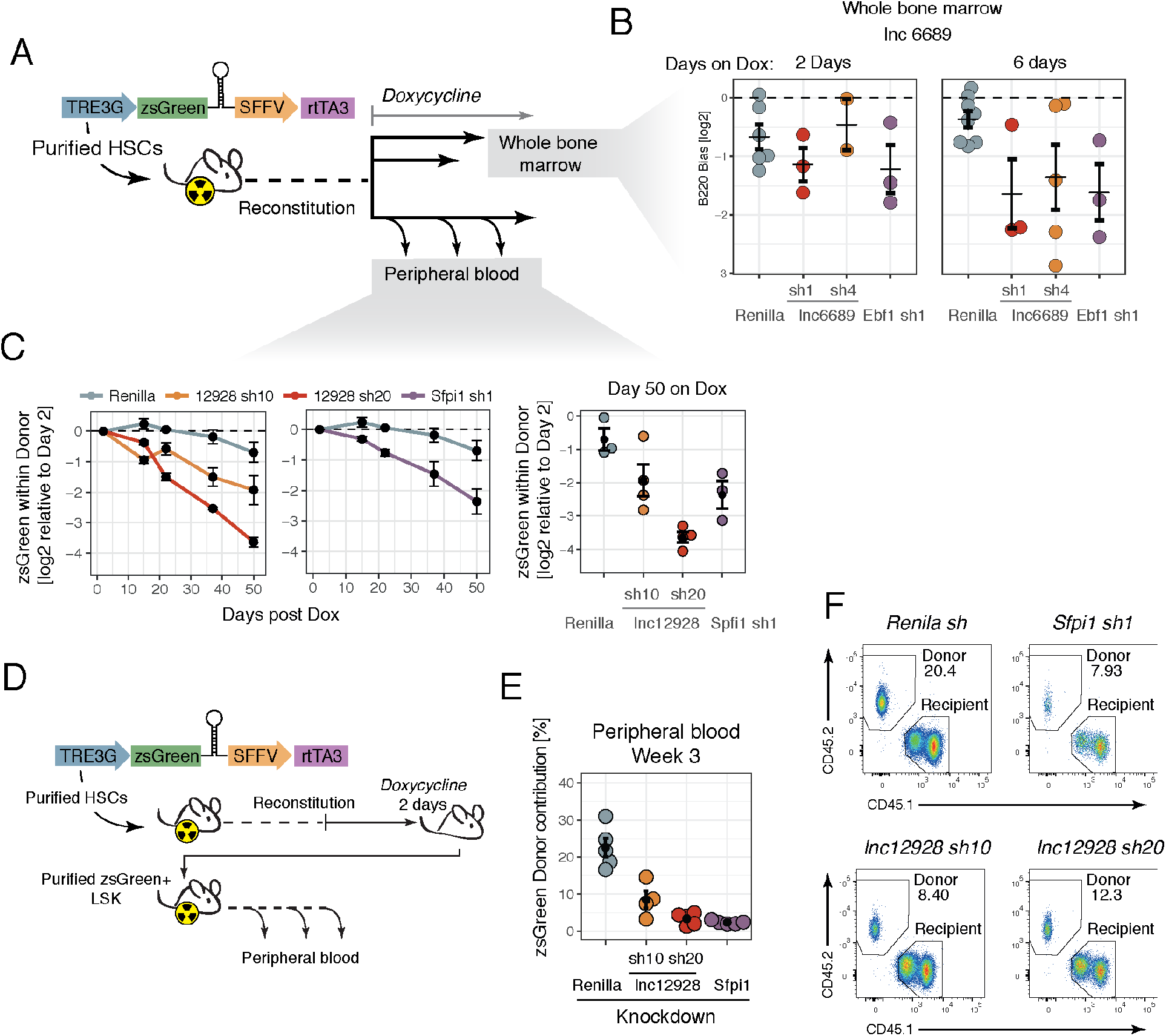
lncRNA 6689 knock down results in B cell depletion and myeloid enrichment, while lncRNA 12928 is required for multi-lineage differentiation. (A) Schematic representation of the vector used and the experimental design. (B) Lineage-bias values for the B lineage in the bone marrow for the knockdowns indicated in animals induced with doxycycline for 2 or 6 days. (C) Average representation of zsGreen-positive cells within donor relative to 2 days post-doxycycline administration for each short hairpin over time (left 2 panels). For the last time point (50 days) the values for each animal are represented (right panel). (D) Schematic representation of the vector used and the experimental design. (E) zsGreen Donor % for each re-transplanted animal at 3 weeks post-transplant. (F) Representative flow cytometry plots of the peripheral blood for the conditions indicated. Error bars represent s.e.m.

### *Lncl2928 (Spehd)* is required for hematopoiesis during regeneration and in homeostasis post-transplantation

Using the same inducible system described above, we sought to investigate whether the *lnc12928* requirement we noted in animals during reconstitution was also observed if the lncRNA knockdown was induced after the hematopoietic system had recovered following transplantation. Essentially, we wished to separate effects on engraftment versus HSC maintenance. We found that the representation of zsGreen-positive cells within the donor compartment decreased over time in the peripheral blood of animals transplanted with HSCs with *lnc12928* knockdown, compared to the initial percentage two days post-doxycycline administration (Fig. 3C). Strikingly, for sh20 the effect of the depletion is similar to that of positive control knockdown, *Sfpi1 (PU1)*, which is known to be essential in HSCs (Fig. 3C).

This confirms that *lnc12928* is required for hematopoiesis, both in a reconstituting setting as well as for homeostasis post-transplant. This lncRNA is highly enriched in all hematopoietic progenitors, with highest expression in the HSC and GMP cell types, but not in any of the differentiated cell types (Fig. S3A). Consequently, these data together suggest this lncRNA must exert an effect at the level of the stem/progenitor cell types. We therefore named this lncRNA *Spehd* (Stem- and Progenitor-enriched required for hematopoietic differentiation).

### Depletion of *Spehd* results in impaired stem and progenitor contribution to hematopoiesis

We next isolated shRNA-expressing stem cells that we could transplant into recipient animals as the sole source of donor cells. Since we expected that for some of the knockdowns these cells were expected to have a deleterious reconstitution phenotype, we used the same competitive transplantation setup as our initial studies (Fig. 2A), using C57BL/6-CD45.1 as recipients and co-injecting whole bone marrow from CD45.1 littermates. In order to isolate the zsGreen-expressing cells, we kept the primary transplanted animals on doxycycline food for 2 days before bone marrow extraction. The recipient animals were similarly kept on doxycycline food to maintain shRNA expression. Due to the extremely low numbers of HSCs in each animal, and the fact that the zsGreen+ cells are only a subset of the donor compartment, we sorted zsGreen+ LSK (Lin-Sca1+ cKit+), which contains a number of multi potent progenitors, in addition to HSCs (Fig. 3D).

We transplanted 3,000 zsGreen-expressing LSK cells into each animal, 4-5 animals per condition, together with a competitive dose of CD45.1 whole bone marrow. At weeks 3 post-reconstitution, we analyzed the peripheral blood for donor contribution resulting from negative-control *(Renilla), Spehd*, or positive-control *(Sfpi1)* knockdown. Whereas the contribution from the *Renilla*-knockdown cells reached 25% of nucleated peripheral blood, on average across five mice, the ability of LSK cells to reconstitute the animals when depleted of *Spehd* was reduced to around 10% on average, and as little as 5% in some animals (Fig. 3E). Although all the cells in the donor compartment should be zsGreen+, we saw a fraction of the donor being zsGreen-negative (presumably due to silencing of the integrated cassette) (Fig. S3C). This donor-derived zsGreen-negative fraction is especially prominent in those conditions where reconstitution potential is impaired, such as with cells carrying *Spehd* sh20 or *Sfpi1* sh1 – when the cells that managed to silence the transgene could have the greatest advantage (Fig. S3C).

The effects are also seen if we look within each lineage for monocyte-macrophages (Ly6g- Cd11b+) and granulocytes (Ly6g+ Cd11b+) and are apparent already at 3 weeks post-transplant (when the donor cells have not yet contributed to the B compartment) (Fig. S3B,C). These data support an essential role for *Spehd* during hematopoietic stem/progenitor cell self-renewal or differentiation that leads to impaired hematopoiesis when the lncRNA is depleted.

### The common myeloid progenitor compartment shows defects in respiratory pathways when *Spehd* is depleted

To investigate the earliest consequence of *Spehd* knockdown and the cell types most affected by its depletion, we isolated cells from animals that had been reconstituted with shRNA-inducible HSCs and administered doxycycline for 6 days. When we examined genes that were differentially expressed upon lncRNA knockdown, comparing CMPs (with either of the two shRNAs) versus control CMPs, we identified 430 genes consistently down-regulated upon suppression of the lncRNA (DESeq2, adjusted p-value < 0.05). Those genes were strongly enriched for components of the oxidative phosphorylation pathway (as annotated by KEGG and Gene Ontology Biological Process) (Fig. 4A). The signature is driven by 40 genes (Fig. S4A) that are downregulated in CMPs (lncRNA knockdown versus control) but to a much lesser extent or not consistently affected in LSK (a stem and progenitor compartment that includes the HSCs) (Fig. 4B, Fig. S4B).

**Figure 4.**
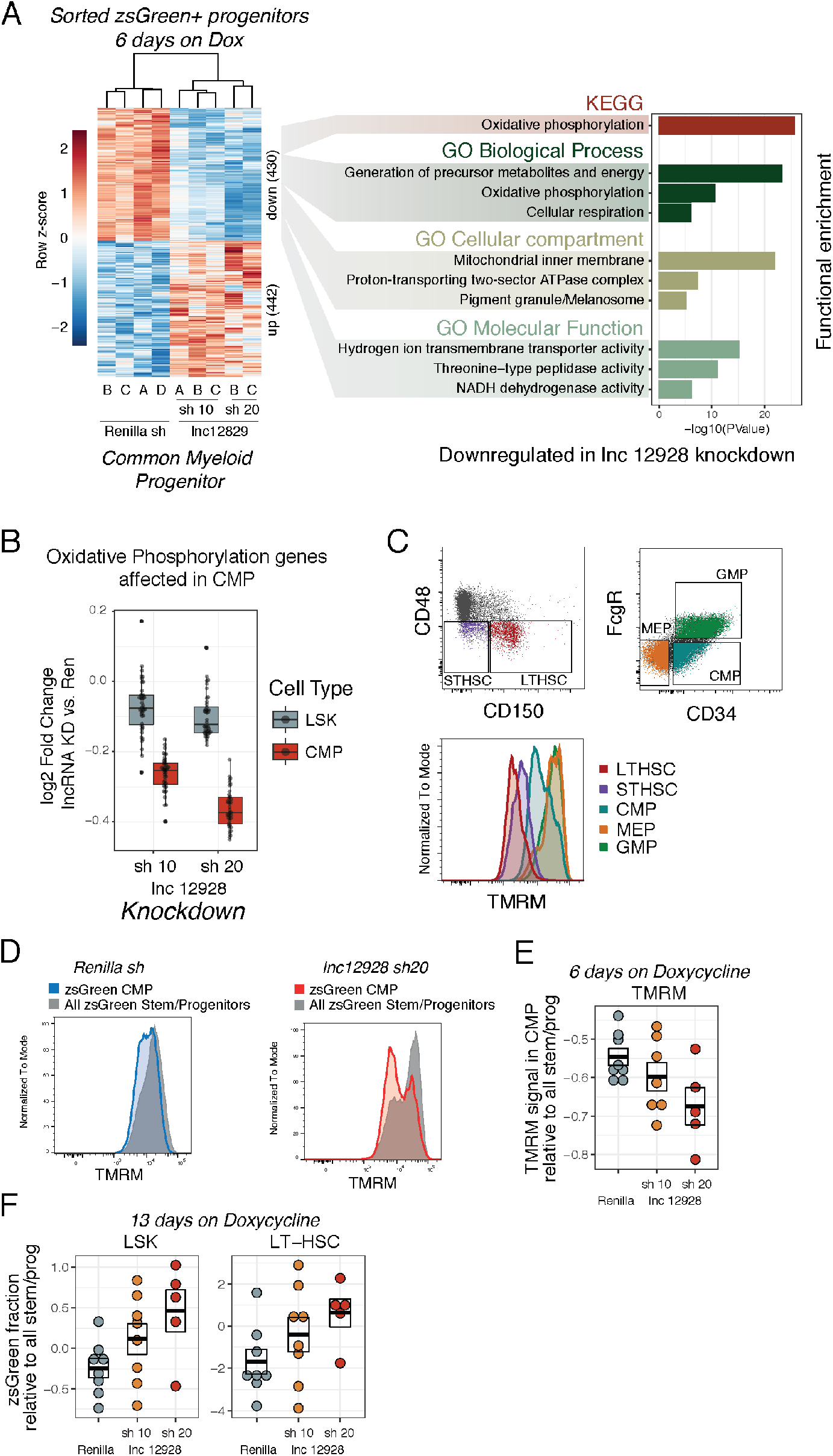
Depletion of lncRNA 12928 results in deficient mitochondrial function in myeloid progenitors. (A) Heat map of all differentially expressed genes (FDR <0.01 DESeq) between lnc12928 knockdown and Renilla in CMP (left) and the results of functional annotation analysis (see Experimental Procedures) (right). (B) Fold change for each short hairpin against the lncRNA relative to Renilla in CMP and LSK for the genes in the oxidative phosphorylation pathway (KEGG) differentially expressed in CMP. (C) Representative flow cytometry plots showing the gating strategy for long term (LT) and short term (ST) repopulating HSC (out of Lin-Sca1+ cKit+) and CMP, GMP, MEP (out of Lin-Sca1-cKi1+) and the corresponding distribution of TMRM intensity for each population. (D) Representative flow cytometry plots of CMP’s TMRM and TMRM in all progenitors and stem cells for Renilla shRNA and short hairpin 20 against lnc12928. (E) TMRM (Geometric mean) in CMP relative to overall TMRM in the sample (in all progenitors and stem cells) per animal. Box represents average and s.e.m. (F) zsGreen in either LSK (left) or LT-HSCs (right) relative to the zsGreen % in all progenitor/stem cells for each animal. Box represents average and s.e.m.

Hematopoietic stem cells are reported to primarily use glycolysis (Takubo et al., 2013) and have lower respiratory capacity than multipotent progenitors or committed progenitors (de Almeida et al., 2017). Moreover, several mutants with mitochondrial and respiration defects have profound hematopoietic defects (Maryanovich et al., 2015; Yu et al., 2013) and HSCs with lower mitochondrial activity have greater reconstitution capacity (Vannini et al., 2016). Therefore, it seemed possible that cells with reduced expression of genes encoding respiratory chain components could encounter a roadblock when their metabolic requirements increase. One such requirement is when HSCs leave their quiescent state and undergo the metabolic awakening that is associated with their commitment to differentiation and expansion.

We therefore asked whether the 40 oxidative phosphorylation-pathway genes affected in CMP depleted of *Spehd* were showed increases in expression during differentiation between LSK and CMP. In the control case *(Renilla* shRNA), all of the genes show, on average, increased expression, and 29 of the 40 are affected consistently, resulting in a statistically significant up-regulation (DESeq2, adjusted p-value < 0.05) (Fig. S4B). Consistent with our previous analysis, these genes fail to become induced in the CMPs depleted of lncRNA *Spehd*, with either sh10 or sh 20 (Fig. S4B).

Considered together, these results indicate a defect in the committed myeloid progenitor population, which fails to activate the expression of genes involved in the oxidative phosphorylation pathway. Defects in this pathway could lead to a deficient metabolic state that could explain the substantial reduction in differentiated cell output we observed in this lncRNA knockdown. Interestingly, *Spehd* was primarily cytoplasmic in leukemia cells (Fig. S4C-E). While the exact mechanism by which *Spehd* exerts its function remains to be elucidated, its localization would seem to exclude direct effects at the transcriptional level.

To further investigate this phenotype, we transplanted animals following the same experimental design used for transcriptomic profiling (Fig. 4A). We tested whether mitochondria function was affected by lncRNA knockdown in the common myeloid progenitor using tetramethylrhodamine methyl ester (TMRM). This cell permeable dye is sequestered in active mitochondria and gives a progressively higher readout as you progress down the hematopoietic lineage in a manner that correlates with commitment status (Vannini et al., 2016) (Fig. 4C). When we measured the level of TMRM for CMPs (calculated as normalized geometric mean relative to the TMRM level of all the stem and progenitor cells – LSK and Lin Sca1-cKit+ together), we saw a reduction in TMRM signal for both short hairpins targeting *Spehd* (Fig. 4D-E). While both short hairpins show a reduction, short hairpin 20 provokes, once again, a stronger phenotype, correlated with its greater potency. These data lend further support to the hypothesis that lncRNA *Spehd* is required in committed progenitors and its depletion causes a deficiency in the metabolic proficiency of these cells. We note TMRM accumulation could also correlate with bulk mitochondrial mass rather than their metabolic capacity.

Notably, when we analyzed animals that have been treated with doxycycline continuously for 13 days, we observed that the relative representation of zsGreen-positive cells within stem-progenitor compartment (LSK)—and specifically within long term repopulating HSCs—was, on average, increased when we knocked down *Spehd* relative to the *Renilla* control (Fig. 4F). This might indicate that stem cells depleted of *Spehd* accumulate after a short-term acute *Spehd* knockdown. Although we did not observe a corresponding decrease in CMPs or other committed progenitors at this time point, these data could support a model in which lncRNA *Spehd* affects the differentiation capacity of hematopoietic stem cells and decreases the respiratory capabilities of committed progenitor cells, which have to rapidly amplify to repopulate the system upon transplantation. We hypothesize that the ability of these defective progenitors to differentiate is reduced, leading to the strong phenotype we consistently observed. While we have only detected an oxidative phosphorylation deficiency in the myeloid progenitors, we have observed that B cells were also greatly reduced when this lncRNA was knocked down. We suspect a similar phenotype could be occurring in the lymphoid compartment at an unexplored time point, although an alternative mechanism that explains the lymphoid deficiency could also be contemplated.

## Discussion

We have previously cataloged lncRNA expression during mouse hematopoiesis and carried out a functional analysis of lncRNAs in the hematopoietic compartment in mouse leukemias as a proof-of-concept (Delás et al., 2017). Here, we have further explored lncRNA function in hematopoiesis, developing a strategy that allowed us to identify lncRNAs that play a role in hematopoietic stem cell self-renewal or differentiation. Studies of hematopoietic stem cells pose many challenges because of their very low abundance and the lack of culture conditions suitable for their expansion. This prompted us to attempt to examine lncRNA function using *in vivo* HSC reconstitutions. While a powerful approach, the number of candidates that can be assessed using such assays is much lower than one can survey in vitro.

For these reasons, we developed a candidate selection strategy based on combining mouse lncRNA expression data, synteny, and human expression data to enrich for lncRNAs with the greatest potential for functional roles in hematopoietic stem and progenitor cells. Our *in vivo* functional assay has revealed two lncRNA with phenotypes in hematopoietic differentiation. We further investigated the phenotype and potential functional relevance of *Spehd*, an RNA expressed in all progenitors and in hematopoietic stem cells and required for hematopoietic differentiation.

The strategy that we developed for candidate selection narrowed our study to 5 lncRNAs and was aimed at identifying non-coding species that could regulate the first steps of HSC commitment into myeloid or lymphoid lineages. However, this combination of differential expression, synteny, and conserved expression is broadly applicable to either other cell types in the hematopoietic lineage or other tissues.

Within the final candidate list, we noted that many of the genes surrounding the lncRNAs of interest had known roles in hematopoiesis (*Cdk6* and *Gata2* for example). Although a conserved role in *cis* cannot be excluded, we suspect these coding and non-coding genes are simply regulated by the same tissue specific elements in both mouse and human, which allows them to fulfill criteria that we set for candidate selection. One major drawback of our selection strategy was that we could have missed a functionally conserved lncRNA that does not show synteny between mouse and human.

*Spehd* emerged from this approached as a lncRNA required for stem cell differentiation. Hematopoietic stem cells with reduced levels of *Spehd* show a decreased capacity for multi-lineage differentiation over time. When Spehd-depleted HSPCs (LSK compartment) were transplanted in a competitive fashion, we saw that their ability to contribute to a number of blood lineages was greatly reduced, indicating either a HSC defect or a broad effect on all progenitors.

Transcriptomic profiling of hematopoietic progenitors after acute induction of two independent shRNAs against *Spehd* revealed reduced expression of genes involved in oxidative phosphorylation in the common myeloid progenitor population. Moreover, the relative levels of TMRM in these same cells was reduced in the presence of this lncRNA knockdown, supporting a deficiency in mitochondria function or a reduction in mitochondrial mass. Even a slight defect in this pathway has the potential to be detrimental during a process where cells exit a quiescent state and have to undergo a rapid expansion, giving rise to a myriad of differentiated cell types. Intriguingly, a recent report has identified the lncRNA here annotated as *lnc6689 (2810468N07Rik)* also as a regulator of oxidative phosphorylation in a microRNA-dependent manner (Sirey et al., 2018). Of course, the major questions that remain are a challenge in lncRNA biology more broadly, namely by what mechanism precisely does this non-coding RNA regulate the differentiation potential and the metabolic capacity of HSCs.

## Experimental Procedures

### Animal Work

Bone marrow transplantations of modified HSCs and subsequent analysis of peripheral blood and bone marrow from these mice were performed at the Cancer Research UK Cambridge Institute (Cambridge, UK). C57BL6J and C57BL/6-CD45.1 were purchased from Charles River (Kent, England) and used at 9–12 weeks old. C57BL/6-CD45.1; cKit-w41 animals used in inducible transplantations were re-derived from embryos provided by David Kent at the Cambridge Institute for Medical Research and transplanted at the same age. Induction of shRNAs was performed by administering doxycycline-containing food (625 mg/kg) for the days indicated at 10 weeks (when bone marrow was analyzed) or 12-13 weeks (when peripheral blood was analyzed) post-transplant. These animal procedures were conducted in accordance with project and personal licenses issued under the United Kingdom Animals (Scientific Procedures) Act, 1986.

### lncRNA expression, human synteny and correlation analysis for candidate selection

The mouse RNAseq libraries from progenitors and differentiated cell types were previously published (Delás et al., 2017). Differential expression among hematopoietic progenitors was analyzed using DESeq2 (Love et al., 2014) and lncRNAs were considered to be differentially expressed if the fold change exceeded 2 and FDR<0.05 in the comparisons indicated in the main text. Synteny was determined by converting the genomic interval for the mouse lncRNAs of interest from the mm10 to hg38 assembly using the UCSC’s liftover tool. The presence of human lncRNAs in these regions was verified using “BEDTools Intersect” with the GENCODE v22 lncRNA annotation. Human RNAseq expression data from cord progenitor types was obtained from previously existing datasets (Chen et al., 2014). Reads were mapped with the STAR aligner (Dobin et al., 2013) against the hg38 assembly, and fragment counting was performed with htseq-count (Anders et al., 2015). To compute expression correlation between mouse and human we first selected lncRNAs with an average expression across the human cell types of more than 20 normalized counts (value determined from the empirical count distributions). DESeq2 was used to calculate variance-stabilized data for each cell type in human and mouse. Expression correlation was computed between the median values for HSC, CMP, GMP and CLP in mouse versus human using Pearson correlation.

### shRNA design and cloning

shRNAs were predicted using the shERWOOD computation algorithm (Knott et al., 2014) as previously described for lncRNAs (Delás et al., 2017). shRNAs were cloned into the appropriate vectors, with ultramiR backbone: ZIP-Neo (constitutive bone marrow transplantations), or T3G-zsGreen-ultramiR-SFFV-rtTA (L3zUSR) (clonal inducible cell lines, inducible bone marrow transplantations) as previously described (Knott et al., 2014).

### Virus production

In brief, virus was prepared in 15 cm dishes using 293FT cells (Thermo Fisher Scientific). The transfection mixture contained 32 μg of DNA vector, 12.5 μg of pMDL, 6.25 μg of CMV-Rev, 9 μg of VSV-G, 200 μg of Pasha siRNA, 125 μl 2.5M of CaCl_2_ brought to 1250 μl with H_2_O and bubbled into 1250 μl 2X HBS. Media was changed to IMDM supplemented with 10% heat-inactivated FBS right before transfection and collected in 16 ml of the same media. 38 ml of viral supernatant was ultracentrifuged for 2.5 hours at 25,000 rpm at 4°C, and resuspended in 100 μl of D-PBS (Gibco). Viral titer was determined by infection of 293FT cells (Thermo Fisher Scientific) at various viral dilutions and percent infection was measured by flow cytometry analysis of the fluorescent protein expressed.

### Isolation, infection, and short-term culture and transplantation of HSCs

Bone marrow from C57BL/6 mice was extracted by flushing, filtered through a 0.30 μm filter and lineage depleted (Mouse Lineage depletion kit, Miltenyi Biotec 130-090-858). Cells were stained with EPCR-PE, CD45-APC, CD150-PE/Cy7 and CD48-FITC. DAPI or LIVE/DEAD Fixable Violet Dead Cell Stain Kit (Thermo Scientific L34963) was used for dead cell exclusion. Sorting of highly pure E-SLAM HSCs (on a FACSAria IIU, BD Biosciences) and short-term (~ 20 hour) culture was performed as previously described (Kent et al., 2009). In short, 1000 live EPCR+CD45+CD150+CD48-lineage negative cells were sorted in 100 μl of media: Iscove modified Dulbecco medium supplemented with 10 mg/mL bovine serum albumin, 10 μg/mL insulin, and 200 g/mL transferrin, 100 U/mL penicillin, 100 μg/mL streptomycin (purchased as BIT from StemCell Technologies), and 10^−4^ M β–mercaptoethanol (Gibco) plus 20 ng/mL interleukin-11 (IL-11; R&D Systems) and 300 ng/mL Steel factor (R&D Systems)). Ultracentrifuged viral supernatant was added aiming for a final concentration of ~2 x10^7^ IU/ml following the sorting. Each well was used to inject four animals; cells were washed prior to injecting to remove remaining viral particles.

Transplants with the constitutive vector (Fig. 2) were performed into lethally irradiated (900 cGy in two split doses) C57BL/6-CD45.1 animals co-injecting 2×10^5^ nucleated whole bone marrow cells (from C57BL/6-CD45.1 animals) per animal. Transplants with the inducible vector (Fig. 3A-C, Fig. 4) were performed into sub-lethally irradiated (450 cGy) C57BL/6-CD45.1; cKit-w41 animals without competitive cells. Re-transplants (Fig. 3C-F) were performed following the constitutive vector protocol (into C57BL/6-CD45.1with competitive dose) using Lin-Sca1+cKit+ (LSK) zsGreen+ cells isolated from transplanted animals. In the case of the re-transplants (Fig. 3E-F), the cells were reinjected the same day they were extracted.

### Peripheral blood extraction and flow cytometry analysis

Blood from transplants with the constitutive vector was analyzed starting from 4 weeks after transplantation and every 4 weeks thereafter. 50–75 μl of blood was extracted from the animals’ tail vein into heparin coated capillary tubes. Red blood cell lysis was performed using Ammonium Chloride Solution (Stem Cell Technologies). Samples were then stained with B220-APC, CD3-AF700, CD11b (Mac1)-APC-Cy7, Ly6G-APC-Cy7, CD45.1-PE, and CD45.2-BV421, and analysis was done on a LSR Fortessa (BD Biosciences). Flow data analysis was performed using FlowJo and statistical analysis using R Studio.

For blood analysis from the inducible vector transplants and re-transplanted animals, blood was stained with CD45.1-AlexaFluor700, CD45.2-BV421, CD45R/B220-APC, Ly-6G-PE, CD11b (Mac-1)-PE-Cy7 and Fixable Viability Dye eFluor™ 780 (eBioscience) for dead cell exclusion. Animals were analyzed from 2 days after doxycycline administration (inducible vector transplants) or at 3 weeks post re-transplant, as indicated in the Figures.

### Bone marrow extraction and flow cytometry analysis

Bone marrow was analyzed from the femurs of euthanized animals at various time points following transplantation. Whole bone marrow was extracted by flushing and was filtered through a 0.3 μm filter. To obtain progenitor populations, 5 × 10^7^ bone marrow cells were lineage depleted (Mouse Lineage depletion kit, Miltenyi Biotec 130-090-858). Cells were stained with FcγR-BUV395, CD34-Biotin-Streptavidin-PE, CD45.1-AF700, CD45.2-V500 or BV510, cKit-APC, Flt3-BV421, IL7Rα-PE-Cy7, Sca1-BV605. Fixable Viability Dye eFluor780 (eBioscience) was used for dead cell exclusion, and analysis was performed on a FACSAria IIU (BD Biosciences). For analysis of differentiated populations, red blood cell lysis was performed on whole bone marrow using ACK Lysing Buffer (Thermo Fisher Scientific). Remaining cells were then stained with B220-APC, CD3-AF700, CD11b (Mac1)-APC-Cy7, Ly6G-APC-Cy7, CD45.1-PE, and CD45.2-BV421, and analysis was done on a FACSAria IIU (BD Biosciences). For Tetramethylrhodamine, Methyl Ester, Perchlorate (TMRM, Thermo Fisher Scientific, Cat no: T668) analysis, freshly isolated bone marrow was incubated in TMRM (20 nM) and Verapamil (50 μM) for exactly 30 min at 37°C, then washed and depleted of differentiated cell types as described above. Lineage depleted cells were stained with Ly-6A/E(Sca-1)-PerCPCy5.5, CD117(cKit)-APC, CD34-AF700, CD16/32 (FcγR)-BUV395, CD48-BV421,CD150-PE-Cy7 and analyzed in a FACSARIA II U (BD Bioscience).

### Cell culture maintenance

MLL-AF9;NRAS^G12D^ AML cells were obtained from the Lowe laboratory (Zuber et al., 2011a) and cultured in RPMI-1640 with GlutaMax (Gibco), supplemented with 10% heat-inactivated FBS (Gibco) and 1% Penicillin/Streptomycin (Gibco) under 7.5% CO2 culture conditions. The cell line established from this model is also known as RN2. A20 cells (purchased from ATCC) were cultured in RPMI-1640 ATCC modification (Gibco), supplemented with 10% heat-inactivated FBS (Gibco), 1% Penicillin/Streptomycin (Gibco), and 0.05 mM β–mercaptoethanol (Gibco). 293FT (purchased from Thermo Fisher Scientific) were cultured as per manufacturer’s instructions. All cell lines tested negative for mycoplasma contamination by RNA-capture ELISA.

### Relative lncRNA quantification by RT-qPCR

Analysis of knockdown efficiency was performed after inducing shRNA expression for 2 days. For all lncRNAs knockdown efficiency was assessed in MLL-AF9;NRAS^G12D^ AML cells (RN2), with the exception of lnc6689 which was assessed in A20 cells. RNA was extracted using the RNeasy Mini Kit (Qiagen), including treatment with the DNase Set (Qiagen). Reverse transcription was performed using Superscript III (ThermoFisher Scientific), with 4 μg of RNA and 1 μl of 50 μM oligo(dT)20. Primers were designed using IDT PrimerQuest tool or chosen from IDT’s pre-designed set when available. Fast SYBR Green (ThermoFisher Scientific) was used for qPCR. Primer pair efficiency was assessed using serial dilutions of cDNA from untreated RN2 or A20 cells, and melting curves were examined to ensure the presence of only one amplicon. *Gapdh* was used as a housekeeping normalization control in the delta-delta-Ct analysis.

### Subcellular Fractionation

Subcellular fractionation of MLL-AF9;NRAS^G12D^ AML cells (RN2) was performed as previously publishes (Gagnon et al., 2014). In brief, 2×10^7^ cells were split in two equal aliquots, one for total RNA isolation and one for cellular fractionation. For fractionation, cells were washed in ice-cold 1X PBS and incubated on ice for 10 min in 380 μl of ice-cold Hypotonic Lysis Buffer, HLB, (10 mM Tris pH 7.5, 10 mM NaCl, 3 mM MgCl_2_, 0.3% Igepal CA-630 (SIGMA-ALDRICH) and 10% glycerol) supplemented with 100 U of SUPERase-In RNase Inhibitor (ThermoFisher Scientific). The sample was centrifuged at 1,000g at 4 °C for 3 min. The supernatant (cytoplasmic fraction) was mixed with 1 mL of RNA Precipitation Solution, RPS, (0.5 mL 3 M sodium acetate pH 5.5 and 9.5 mL ethanol) and stored at −20°C for at least 1 hour. The pellet (semi-pure nuclei) was washed three times with 1 mL of ice-cold HBL and resuspended in 1 mL of Trizol LS (ThermoFisher Scientific). The cytoplasmic fractions (in RPS) was vortexed for 30 s and then centrifuged at 18,000g at 4 °C for 15 min. The pellet was washed in ice-cold 70% (vol/vol) and resuspended in 1 mL of Trizol LS. RNA from either fraction or from the total sample (in Trizol) was extracted by adding 200 μl of Chloroform (SIGMA-ALDRICH), mixing and incubating at room temperature for 10 min. The phases were subsequently separated by centrifugation at 18,000g at room temperature for 10 min. An equal volume of 70% (vol/vol) ethanol was added to the aqueous phase and RNA was isolated using the RNeasy Mini Kit (Qiagen), including DNase treatment (Qiagen), performed according to manufacturer’s instructions. The RNA was eluted in 30 μl of Nuclease-free water. Reverse transcription was performed using SuperScriptIII (ThermoFisher Scientific) as described above but using equal volumes of RNA solution from each fraction sample (nuclear, cytoplasmid or total RNA). Two primer pairs were used for lnc12928, in addition to primers for *Malat1* as a nuclear control and *Gapdh* as a cytoplasmic control. Cytoplasmic enrichment for each primer pair was calculated as the 2^-(Cytoplasmic Ct – Total Ct)^. The nuclear enrichment was calculated accordingly. This does not provide an absolute measurement of the proportion of lncRNA in each fraction but rather allows one to compare the enrichment in each fraction the known controls.

### Single molecule RNA FISH

SmRNA FISH probes were ordered from *Stellaris* conjugated to Quasar^®^ 570 Dye. Malat1 and Gapdh mouse controls were selected from the ShipReady *Stellaris* probe sets. The protocol was performed according to manufacturer’s instructions for cells in suspension (Biosearch Technologies). 5 × 10^6^ cells were washed in 1 mL of 1X PBS and the pellet was fixed in 1 mL of Fixation Buffer (3.7% formaldehyde in 1X PBS) at room temperature for 10 min. The fixed cells were washed three times with 1X PBS and then permeabilized at 4°C for at least 1 hour using 70% (vol/vol) ethanol. 500 μl of fixed/permeabilized cells were washed in 500 μl of Wash Buffer A (10% formamide in 1X Stellaris Wash Buffer A) before incubating in 100 μl of Hybridization Buffer (10% formamide in Stellaris Hybridization Buffer) containing 125 nM probe at 37°C in the dark overnight. The sample was centrifuged to pellet the cells and 50% of the Hybridization Buffer was removed. The pellet was then washed in Wash Buffer A and incubated in the dark at 37°C for 30 min with 500 μl of this same buffer. The nuclei were stained with NucBlue Fixed Cell Stain (ThermoFisher Scientific), the cells were washed with Stellaris Wash Buffer B and seeded on a clean glass microscope slide in one drop of ProLong Diamond Antifade Mountant (ThermoFisher Scientific). The cells were imaged in a Nikon TE2000 Widefiled inverted microscope. Z-stacks were acquired by sampling every 0.3 μm. For subcellular localization analysis of the lnc12928 probe, the nuclear or cytoplasmic localization or the signal was established in the Z stacks where the smFISH was detected. The fraction of cytoplasmic or nuclear signal per cell was calculated.

### RNA-seq from lncRNA-depleted bone marrow progenitor populations and data analysis

LSK, CLP, CMP, and GMP populations were sorted from the bone marrow isolated from transplanted mice as described above. RNAseq libraries were prepared as previously described (Delás et al., 2017). In brief, RNA was extracted using the NucleoSpin RNA XS kit (Machery Nagel). cDNA and libraries were prepared used the SMART-Seq v4 Ultra Low Input RNA Kit for Sequencing (Clontech), SeqAmp DNA Polymerase (Clontech), and Low Input Library Prep Kit (Clontech). Samples were pooled and run on a HiSeq 4000.

RNAseq libraries were mapped with STAR aligner (Dobin et al., 2013) against the mm10 mouse genome assembly using default parameters. Duplicate alignments were removed from the resulting BAM files with Picard (http://broadinstitute.github.io/picard). HTSeq-count (Anders et al., 2015) was used to calculate gene counts and subsequently input them into DESeq2 (Love et al., 2014) for quality control analysis, size normalization and variance dispersion corrections. For functional annotation analysis, DESeq2 was used to calculate differentially expressed genes (FDR<0.1). Significantly downregulated genes in CMPs upon lncRNA knockdown were use as input for DAVID 6.7 (Huang et al., 2009). The categories shown are the result of performing Functional Annotation clustering for each category (Gene Ontology Biological Process, GO Molecular Function, GO Cellular Compartment of KEGG). Terms with Bonferroni-corrected p-value <0.05 for the first 5 clusters are represented.

---

**Fig. S1.**
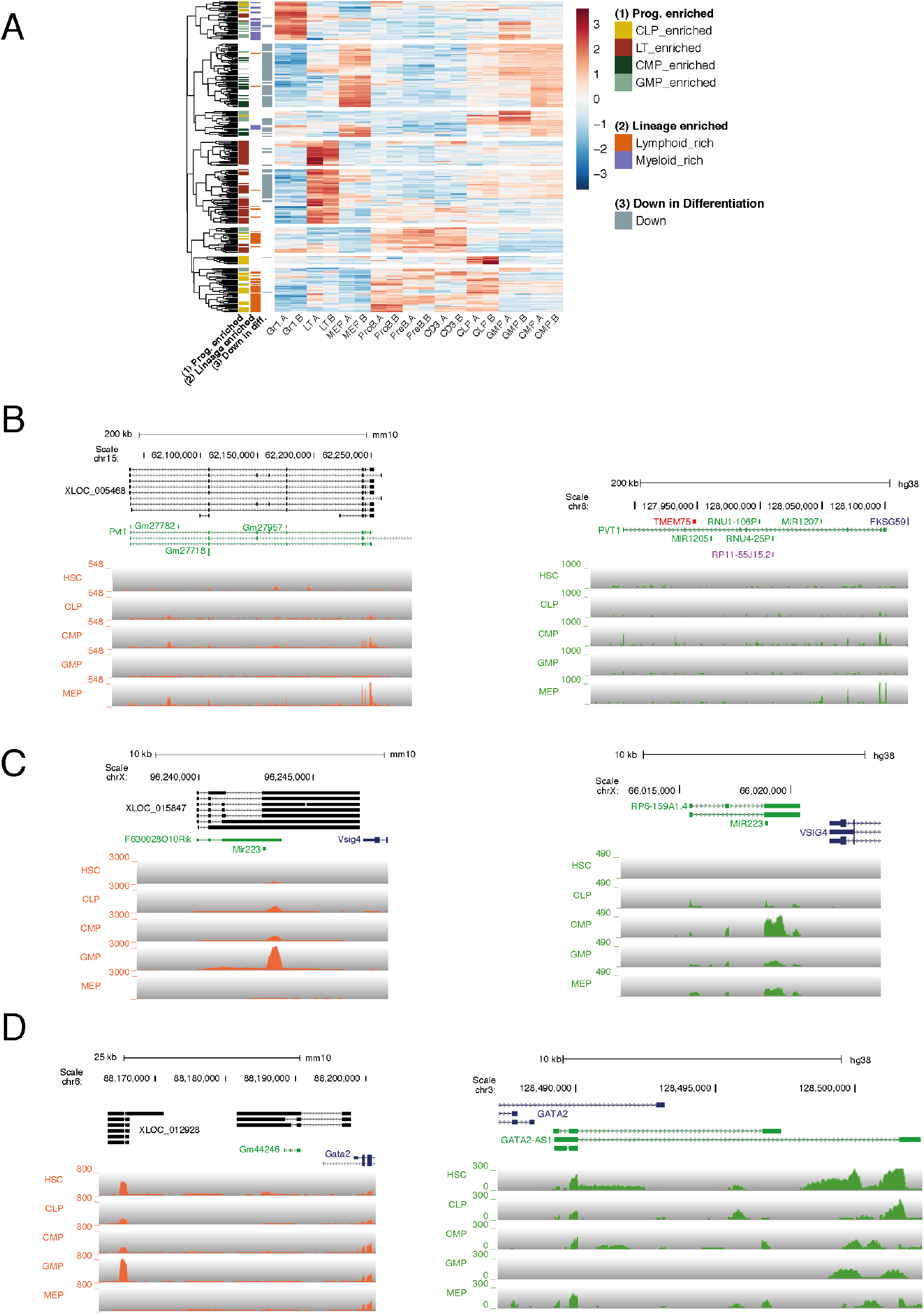
(A) Heat map representing row-scaled expression for all the lncRNAs selected based on their expression patterns (B-D) Genome browser plots of all the remaining 3 lncRNAs selected for *in vivo* studies not shown in Fig.1

**Fig. S2.**
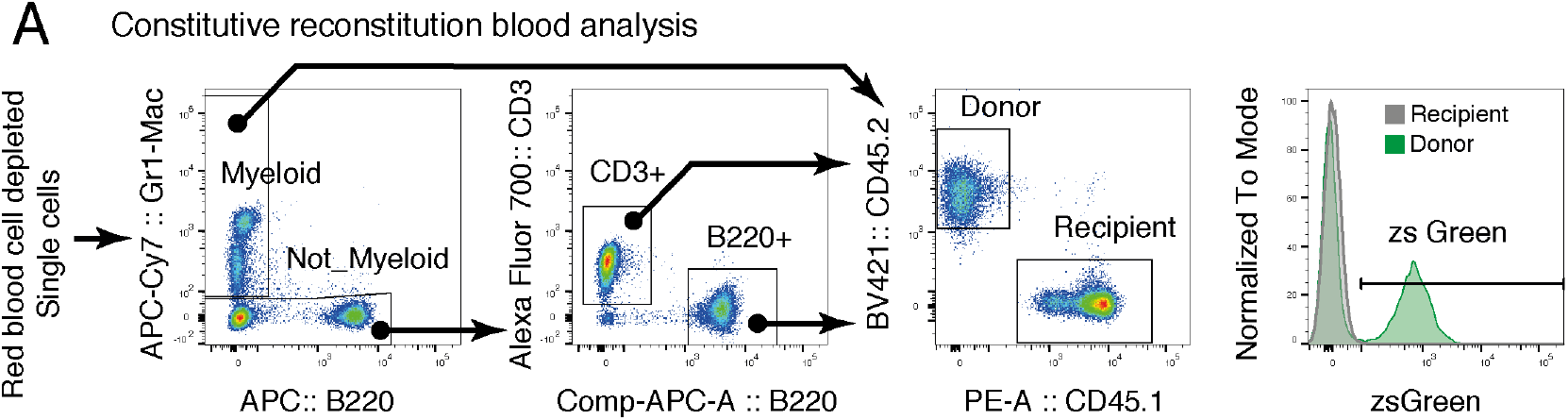
(A) Summary of the flow cytometry analysis performed in the peripheral blood of animals transplanted with HSCs transduced with the constitutive vector.

**Fig. S3.**
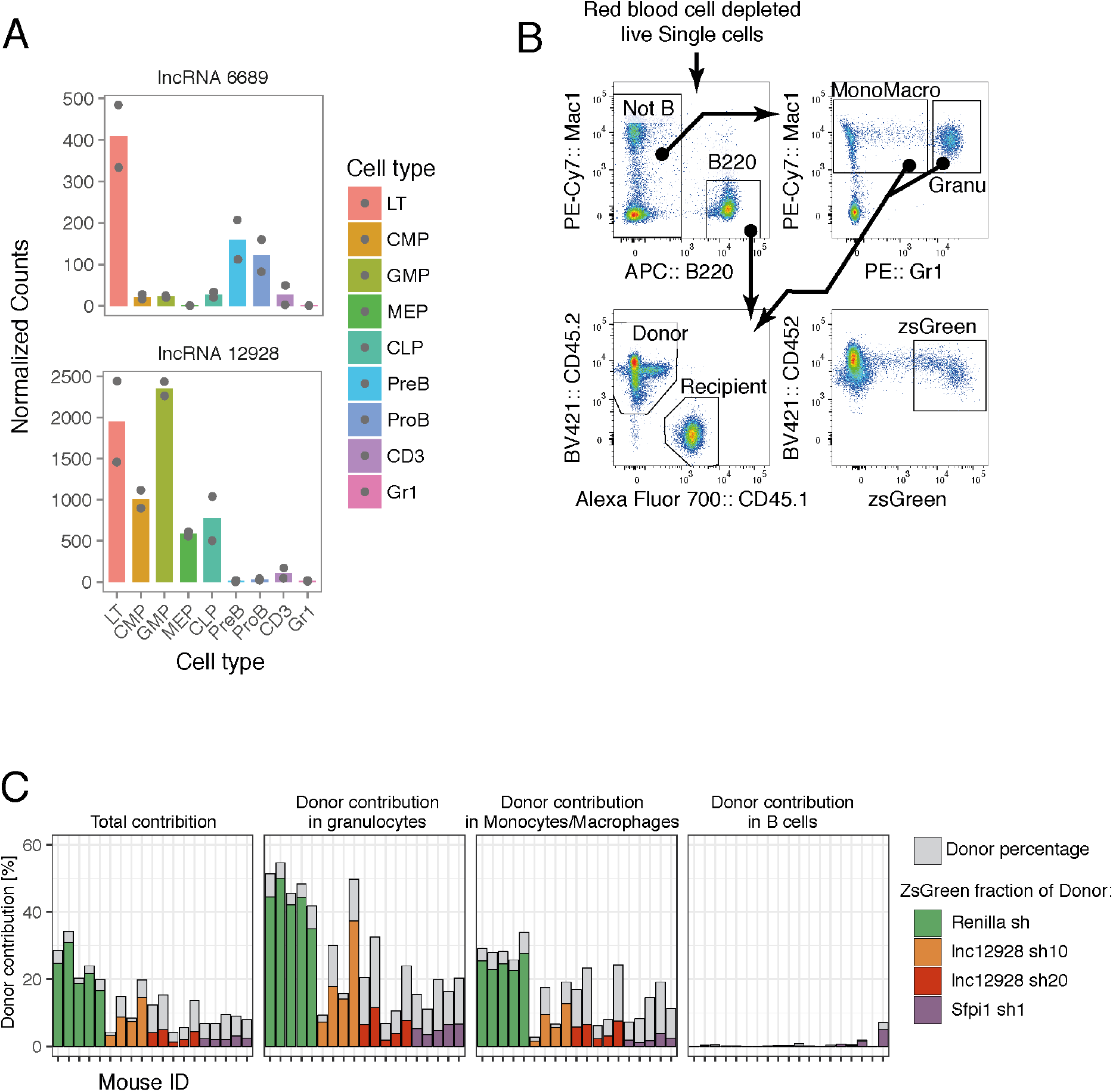
(A) Expression for lnc6689 (upper panel) and lnc12928 (lower panel) in all the cell types analyzed in our previous study. (B) Summary of the ñow cytometry analysis performed in the peripheral blood of animals transplanted with HSCs transduced with the inducible vector. (C) Bar graph representing the overall donor contribution (grey) and the zsGreen+ fraction within it (colored part) for each animal at 3 weeks post-transplant for overall blood or the lineages indicated.

**Fig. S4.**
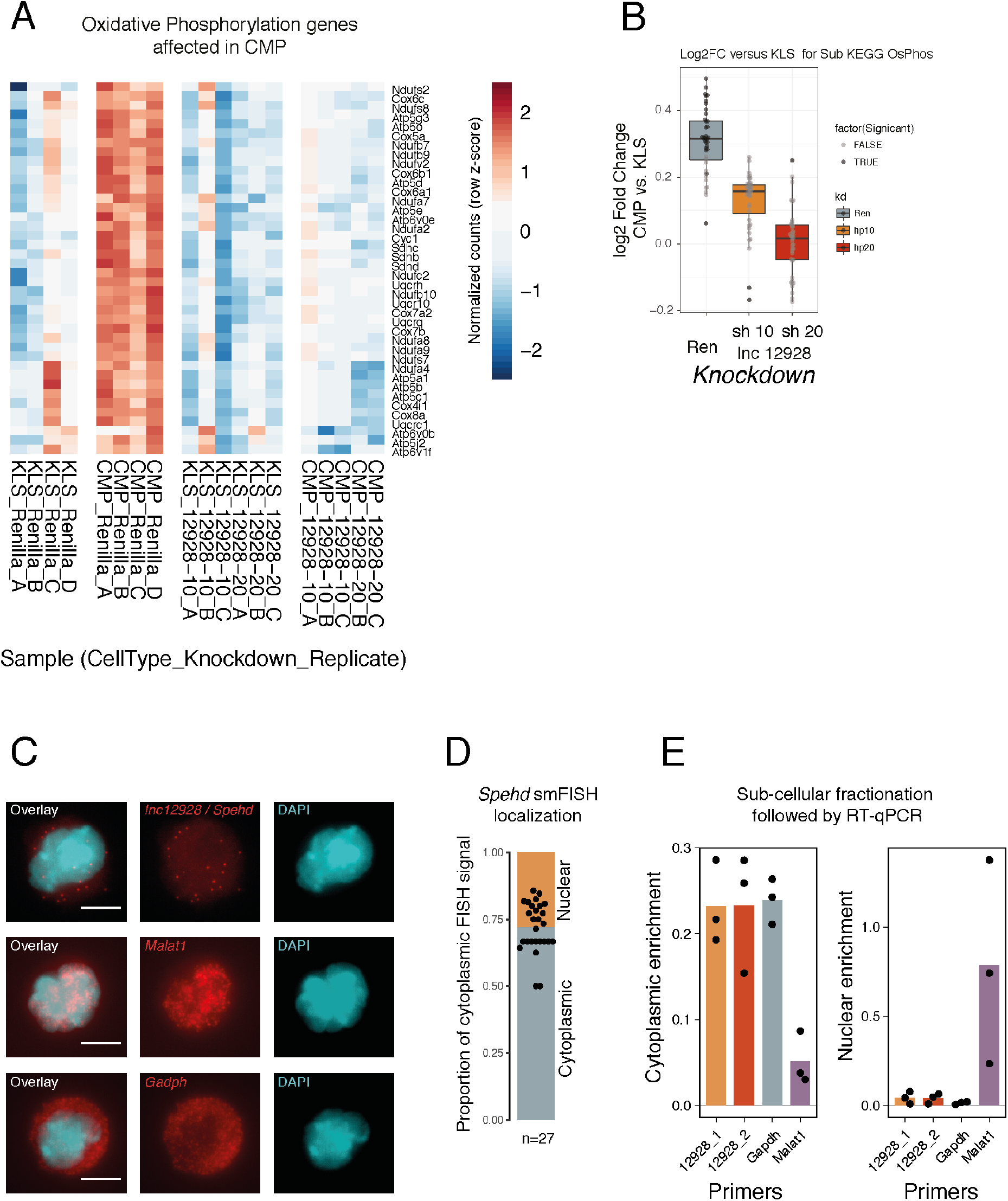
(A) Heat map representing the row-scaled expression in the indicated samples for the 40 oxidative phosphorylation genes (KEGG) affected by lnc12928 knockdown in CMPs. (B) Fold change in expression of the same genes in CMPs compared to LSK for each indicated short hairpin. (C) Representative single molecule FISH (smFISH) images for *lnc12928/Spehd, Malat1* (nuclear control) or *Gapdh* (cytoplasmic control). Images are maximum intensity projections. (D) Quantification of the smFISH spots localization per cell for *lnc12928/Spehd*. Bar height indicates overall average. Each dot is a cell. Cytoplasmic/nuclear localization was determined based on their colocalization with DAPI signal in the stacks where the signal was observed. (E) Cytoplasmic and nuclear enrichment (see Methods) for *lnc12928/Spehd, Malat1* (nuclear control) or *Gapdh* (cytoplasmic control). 12928_1 and 12928_2 correspond to the two primer pairs for lnc12928.

## Tables

Supplementary Table 1: list of *in vivo* lncRNAs with human and mouse coordinates

## Author Contributions

Conceptualization M.J.D and G.J.H; Methodology M.J.D, B.T.J, T.K.; Software S.R.V.K.; Formal Analysis M.J.D., N.E.; Investigation M.J.D, B.T.J., T.K., S.V., E.M.M., S.A.W., E.M.S.; Writing – Original Draft M.J.D., B.T.J; Writing – Review & Editing M.J.D., B.T.J., T.K., G.J.H; Supervision M.J.D, G.J.H; Funding Acquisition G.J.H.

## Acknowledgements

We would like to thank Ilaria Falciatori for helpful discussions and suggestions and Giorgia Battistoni and Cristina Jauset for experimental assistance. We also thank David Kent and Tina Hamilton for the C57BL/6-CD45.1; cKit(w41) mouse line. This work was supported by Cancer Research UK. We specifically thank the Cancer Research UK Cambridge Institute Biological Resource Unit, Flow Cytometry, and Genomics Cores for their support throughout this project. M.J.D was supported by a PhD Fellowship from the Boehringer Ingelheim Fonds. G.J.H. is a Wellcome Trust Investigator, Royal Society Wolfson Research Professor, and was a Howard Hughes Medical Institute Investigator.

